# Stromal ISLR promotes intestinal regeneration and cancer by suppressing epithelial Hippo signaling via FAT1

**DOI:** 10.1101/740472

**Authors:** Jiuzhi Xu, Yang Tang, Xiaole Sheng, Yuhua Tian, Min Deng, Sujuan Du, Cong Lv, Yongli Song, Pengbo Lou, Yongting Luo, Yuan Li, Bing Zhang, Yanmei Chen, Zhanju Liu, Yingzi Cong, Maksim V. Plikus, Qingyong Meng, Zhaocai Zhou, Zhengquan Yu

## Abstract

The YAP signaling activation in epithelial cells is essential for intestinal regeneration and tumorigenesis. However, the molecular mechanism linking stromal signals to YAP-mediated intestinal regeneration and tumorigenesis is poorly defined. Here we report a stroma-epithelia YAP signaling axis essential for stromal cells to modulate epithelial cell growth during intestinal regeneration and tumorigenesis. Specifically, upon inflammation and in cancer, an oncogenic transcription factor ETS1 in stromal cells induces expression of a secreted protein ISLR that can directly binds to a transmembrane protocadherin FAT1 on the surface of epithelial cells. This binding suppressed the Hippo signaling by disrupting MST1-FAT1 association, resulting in YAP signaling activation. Deletion of *Islr* in stromal cells in mice markedly impaired intestinal regeneration, and suppressed tumorigenesis in the colon. Moreover, the expression of stromal cell-specific ISLR and ETS1 significantly increased in inflamed mucosa of human IBD patients and in human colorectal adenocarcinoma, accounting for the epithelial YAP hyperactivation. Collectively, our findings uncovered a molecular mechanism governing signals for communication between stroma and epithelium during tissue regeneration and tumorigenesis.

## Introduction

Inflammatory bowel disease (IBD), which includes ulcerative colitis (UC) and Crohn’s disease (CD), is a type of chronic inflammatory disease associated with increased risk of colorectal carcinoma (CRC). IBD is characterized by submucosal accumulation of inflammatory cells and damage to epithelial cells (Kim & Cheon, 2017, Romano, F et al., 2016). This damage triggers a complex repair program that activates proliferation of epithelial cells that ultimately repopulate the damaged epithelium. When the damage cannot be repaired, the chronic proliferation caused by constant inflammation can drive the development of CRC (Romano et al., 2016). During tissue repair and CRC development, crosstalk between intestinal epithelium and the underlying mesenchyme is critical for promoting epithelial cell proliferation. Mesenchymal stromal cells form the supportive microenvironment that maintains intestinal epithelial architecture during homeostasis (Powell, Pinchuk et al., 2011), become reactive upon inflammation and tissue damage, and further transform into tumor-associated stromal cells upon CRC (Bussard, Mutkus et al., 2016). Reactive or tumor-associated stromal cells can contribute to epithelial cell proliferation in a variety of ways, including through the production of cytokines, such as IL-6 (Francescone, Hou et al., 2015, Zhu, Cheng et al., 2014). Although the importance of mesenchymal stromal cells in promoting epithelial cell proliferation is well established, how stromal signals regulate their proliferation in the context of colitis and CRC remains unclear. A comprehensive understanding of the underlying paracrine mechanisms is of broad interest, because it can uncover new signaling targets for treating these major intestinal diseases.

The Hippo pathway is an evolutionarily conserved signaling cascade, consisting of the core kinase components MST1/2 and LATS1/2, and the transcriptional coactivators YAP and TAZ, and the transcriptional factors TEADs (Huang, Wu et al., 2005, Meng, Moroishi et al., 2016, Pan, 2010). As a tumor suppressor pathway, Hippo signaling is constitutively active under physiological conditions: the MST1/2 kinases phosphorylate and activate LATS1/2 kinases, which in turn phosphorylate YAP/TAZ, leading to cytoplasmic retention and degradation of the latter. However, in the context of tissue regeneration and tumorigenesis, Hippo signaling is frequently dampened, enabling activation of YAP for cell proliferation. Many reports have indicated that YAP/TAZ activity is essential for intestinal epithelial regeneration and tumorigenesis (Cai, Zhang et al., 2010, Gregorieff, Liu et al., 2015, Pan, 2010, Serra, Mayr et al., 2019, Taniguchi, Wu et al., 2015, Yui, Azzolin et al., 2018). Recently, the transmembrane protocadherin FAT1 has been identified as an upstream activator of the Hippo pathway and its mutation is found in many cancers (Ahmed, de Bock et al., 2015, Gee, Sadowski et al., 2016, Li, Razavi et al., 2018, Martin, Degese et al., 2018). It is likely that FAT1 senses extracellular milieu and relays signals to the intracellular Hippo components in epithelial cells. However, the extracellular signal that modulates FAT1 activity during regeneration and tumorigenesis has not been defined, and it also remains unknown how stromal signals regulate epithelial YAP/TAZ activity, especially in colitis and CRC.

ISLR, also known as Meflin, is a conserved protein containing an immunoglobulin (Ig)-like domain and five leucine-rich repeat (LRR) domain (Nagasawa, Kubota et al., 1997, Nagasawa, Kudoh et al., 1999). Recently, ISLR has been identified as a marker of mesenchymal stromal cells, and it functions as a secreted protein to maintain the undifferentiated state of stromal cells (Maeda, Enomoto et al., 2016). Recently, two reports have shown the importance of ISLR in promoting muscle regeneration and cardiac tissue repair (Hara, Kobayashi et al., 2019, Zhang, Zhang et al., 2018). However, it is unknown whether stromal Islr regulates intestinal epithelial regeneration and CRC. Given its abundant expression in stromal cells, it is intriguing to postulate that stroma-secreted ISLR can transduce signals from the microenvironment to epithelial cells, thus mediating stroma-epithelium communications during regeneration and CRC development.

In this study, we utilized stromal cell-specific conditional knockout (cKO) mouse model to demonstrate that stromal cell-secreted ISLR promotes intestinal regeneration and concomitantly acts as a protumorigenic factor in CRC. Mechanistically, extracellular ISLR secreted by stromal cells directly binds to FAT1 located at the surface of epithelial cells, and inhibits its activity, ultimately resulting in Hippo signaling suppression and Yap activation. Thus, we uncovered a previously unrecognized regulatory mechanism governing stroma-epithelium communication essential for tissue regeneration and tumorigenesis.

## Results

### *Islr/ISLR* is increased in stromal cells of the inflamed mucosa in IBD and CRC

To begin elucidating a potential role for ISLR in intestinal homeostasis and in colitis, we first examined its expression patterns in the intestine. Both qRT-PCR and Western blotting assays showed that *Islr* is abundantly expressed in the intestinal lamina propria, but is nearly undetectable in the intestinal epithelium of mice (Fig 1A **and** B). Furthermore, we demonstrate that *Islr* was primarily expressed in non-hematopoietic and non-epithelial cells (CD45^−^Epcam^−^CD90^+^ and CD45^−^Epcam^−^CD90^−^ cells), while it was also mildly expressed in CD45^+^ immune cells and CD45^−^Epcam^+^ epithelial cells (Fig 1C). *In situ* hybridization also showed primary expression pattern of *Islr* in stromal cells of both the small intestine and the colon (Fig 1D). Interestingly, *Islr* levels are the highest at embryonic stage when intestinal epithelium is developing, then decrease to an intermediate level at postnatal day 10; and drop to the lowest level in adulthood (Figs 1D **and EV1A**), suggesting a potential function in promoting cell proliferation.

**Figure 1.**
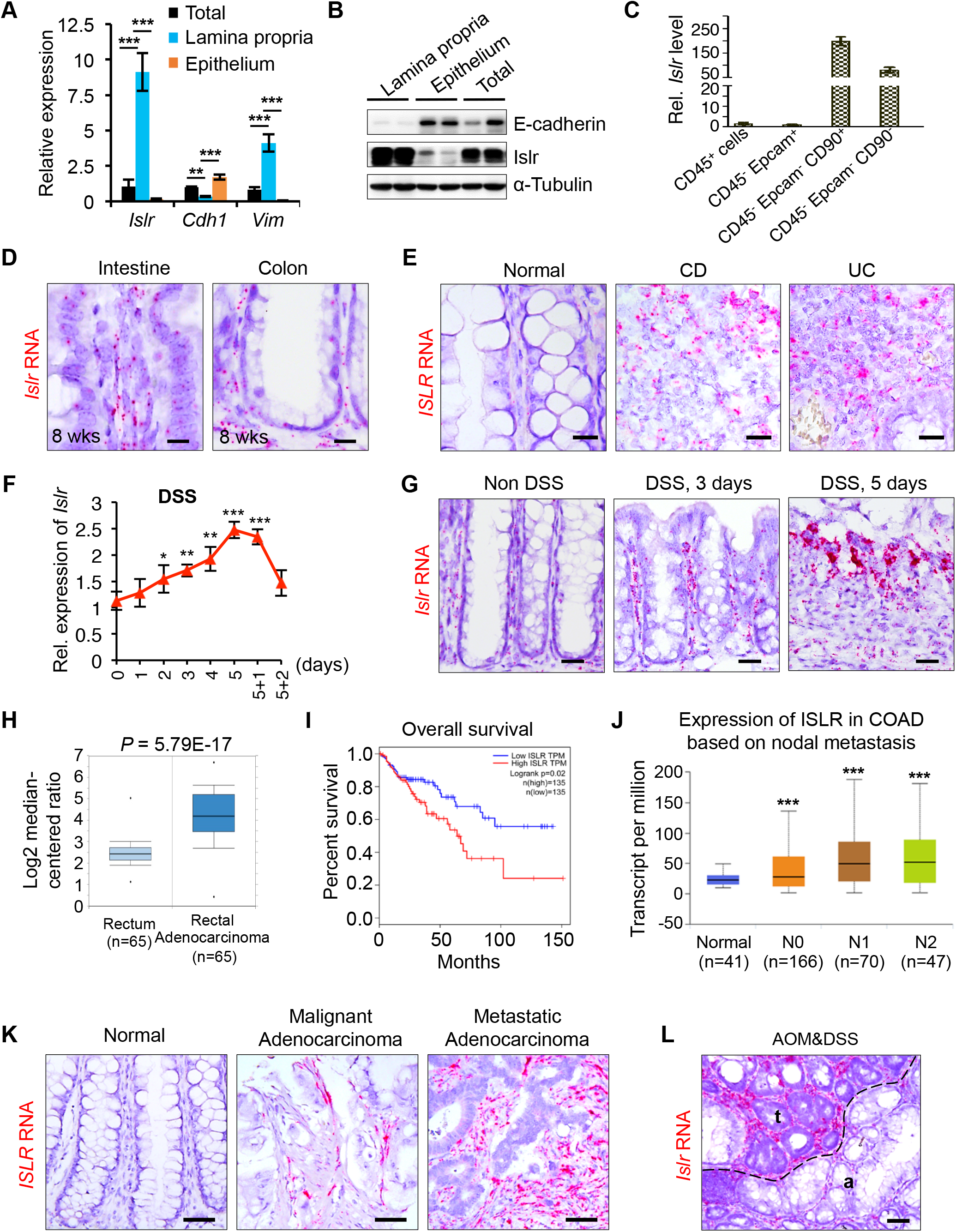
*Islr* is primarily expressed in stromal cells and upregulated in colitis and colorectal cancer. (A) qRT-PCR analysis for *Islr*, *Chd1* and *Vim* in total intestinal tissues, intestinal epithelium and lamina propria. n = 4. \*\**P* < 0.01; \*\*\**P* < 0.001. (B) Western blotting for Islr in total intestinal tissues, intestinal epithelium and lamina propria. α-Tubulin was used as a negative control. (C) qRT-PCR analysis for *Islr* in CD45^+^, CD45^−^Epcam^+^, CD45^−^Epcam^−^CD90^+^ and CD45^−^Epcam^−^CD90^−^ cells sorted from intestinal tissues. n = 3 technical replicates. (D) *In situ* hybridization for *Islr* with RNAscope probe in intestine and colon from 8-week-old mice, showing that *Islr* is primarily expressed in stromal cells. Negative and positive controls were shown in supplementary Figure 1A. Scale bar: 25 μm. (E) *In situ* hybridization for *ISLR* with RNAscope probe in human mucosa, inflamed mucosa from CD and UC patients. Scale bar: 25 μm. (F) qRT-PCR analysis showing the dynamic changes of *Islr* in colonic tissues from mice upon DSS treatment and after DSS removal. n = 4 at each time points. \**P* < 0.05; \*\**P* < 0.01; \*\*\**P* < 0.001. (G) *In situ* hybridization for *Islr* with RNAscope probe in mouse colons without or with 3-day or 5-day DSS treatments. Scale bar: 50 μm. (H) The cancer genome atlas (TCGA) RNA-seq analysis showing that *ISLR* is upregulated in human rectal adenocarcinoma relative to normal rectal tissues. (I) Kaplan-Meier survival curve of 270 CRC cases. *P* = 0.02. (J) *ISLR* level increased with nodal metastasis in colorectal cancer patients. N0, no regional lymph node metastasis; N1, metastases in 1 to 3 axillary lymph nodes; N2, metastases in 4 to 9 axillary lymph nodes; N3, metastases in 10 or more axillary lymph nodes. \*\*\**P* < 0.001. (K and L) *In situ* hybridization for *ISLR/Islr* with RNAscope probes in normal human colon, human malignant adenocarcinoma, metastatic adenocarcinoma (K), and in mouse colon tumors from AOM-DSS model (L). t indicating tumor; a indicating adjacent tissues of tumor. Scale bar: 50 μm. In A and F, data are presented as mean ± SD. \**P* < 0.05; \*\**P* < 0.01; \*\*\**P* < 0.001 (Student’s t-test).

Next, we examined *Islr/ISLR* expression in response to pathological stimuli, including inflammation. *In situ* hybridization showed that *ISLR* levels become dramatically elevated in inflamed mucosa from both CD (n=10) and UC (n= 10) patients, while they were low in unaffected intestinal tissues (Fig 1E). We then induced experimental colitis in mice by administering dextran sulfate sodium (DSS) in drinking water. Consistently, DSS-induced inflammation progressively resulted in marked upregulation of *Islr* in the colon, which gradually recovered to basal levels after DSS removal (Figs 1F **and** G, **and EV1B**).

To further evaluate the relevance of ISLR in inflammation-induced tumorigenesis, we analyzed clinical samples from human CRC patients. *ISLR* was dramatically increased in human CRC tissues relative to normal human colon tissues (Fig 1H), and its expression levels were positively correlated with poor survival and nodal metastasis (Figs 1I **and** J**, and EV1C**). *In situ* hybridization also showed that *ISLR* was primarily expressed in the stromal cells of human colorectal tumors (Fig 1K), and in mouse colon tumors from an azoxymethane (AOM)-DSS model mimicking inflammation-driven colorectal adenocarcinoma (Fig 1L). Taken together, we demonstrate that *Islr/ISLR* is primarily and specifically expressed in stromal cells in the intestine and becomes markedly upregulated in inflamed mucosa and in CRC.

### ETS1 activity induces *Islr* expression in the context of colitis and CRC

To understand how *Islr* expression is regulated, we analyzed the potential binding sites of transcription factors in the 2-kb region upstream from the transcription start site (TSS) of the *Islr* gene locus using the JASPAR database. The analysis identified two ETS1 binding sites (Fig 2A). ETS1 has been identified as an oncogenic transcriptional factor, and its higher expression is associated with higher grading, increased invasion and poorer survival in most cancer types (Dittmer, 2015). Interestingly, *Ets1*/*ETS1* is also primarily expressed in the intestinal stromal cells (Fig 2B **and** C) and is markedly increased in inflamed mucosa from UC and CD patients, experimental colitis and human colorectal adenocarcinoma (Fig 2C-E). The expression levels of ETS1 were positively correlated with ISLR in CRC tissues (Fig 2F). Together, *Ets1*/*EST1* exhibits an overlapping expression pattern with *Islr/ISLR*.

**Figure 2.**
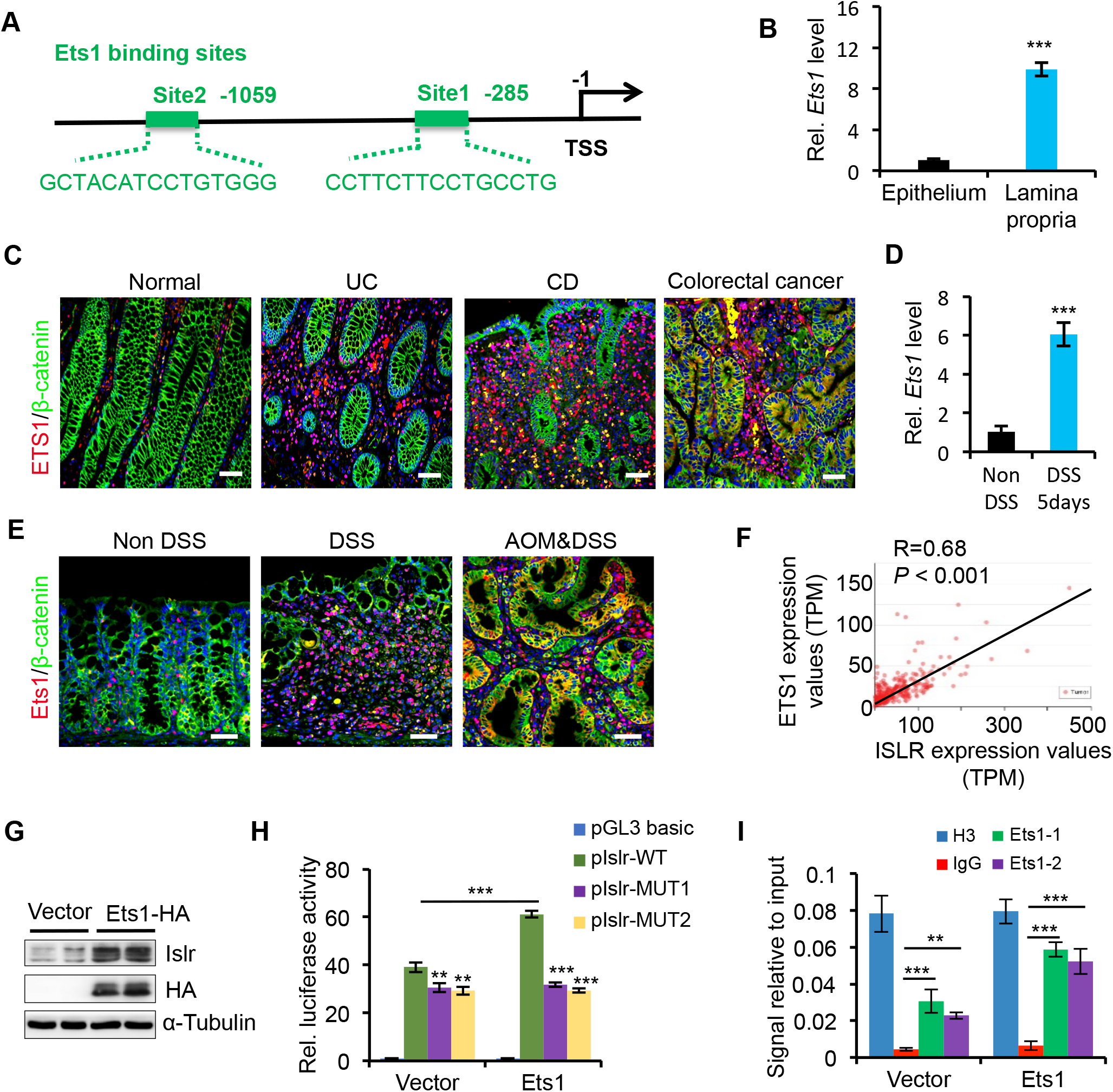
Ets1 activity directly regulated Islr expression. (A) The schematic diagram showing two potential Ets1 binding sites in the *Islr* promoter. (B) qRT-PCR analysis for *Ets1* in colonic epithelium and lamina propria. \*\*\**P* < 0.001. (C) Double immunofluorescence for ETS1 and β-catenin in intestinal biopsies of healthy controls and inflamed mucosa from UC patients. Scale bar: 50 μm. (D) qRT-PCR for *Ets1* in colons with or without 5-day DSS treatment. n = 4. \*\*\**P* < 0.001. (E) Double immunofluorescence for Ets1 and β-catenin in normal mouse colons without DSS treatment, mouse colons with 5-day DSS treatments, and AOM-DSS mouse colon tumors. Scale bar: 50 μm. (F) Pearson correlation analysis on ETS1 and ISLR in colorectal adenocarcinoma TCGA RNA-seq (*P* < 0.001; R = 0.68). (G) Western blotting for ISLR and HA in lysates of UC-MSCs transfected with pCMV5-HA or pCMV5-ETS1. α-Tubulin was used as a loading control. (H) Luciferase activity in lysates of UC-MSCs transfected with luciferase reporter plasmids of pGL3-basic empty vector (basic), wild type *ISLR* promoter or mutant promoter with mutation of ETS1 binding sites under normal and ETS1 overexpression conditions. \*\**P* < 0.01; \*\*\**P* < 0.001. n = 3. (I) Chromatin immunoprecipitation assay was carried out on UC-MSCs cells using antibodies against ETS1 and Histone 3. The antibody against Histone 3 was used as a positive control. IgG was used as a negative control. The enrichment of ETS1 binding to *ISLR* promoter was quantified using qPCR. \*\**P* < 0.01; \*\*\**P* < 0.001. n = 3 technical replicates. In B, D, H and I, data are presented as mean ± SD. \**P* < 0.05; \*\**P* < 0.01; \*\*\**P* < 0.001 (Student’s t-test).

To verify the potential regulatory effect of ETS1 on the expression of *ISLR*, we overexpressed *ETS1* in umbilical cord mesenchymal stem cells (UC-MSCs) and found that *ETS1* overexpression could induce *ISLR* upregulation (Fig 2G). Luciferase reporter assays revealed that ETS1 overexpression was able to activate *ISLR* promoter activity, while mutation in the *ETS1* binding site blocked it (Fig 2H). Furthermore, chromatin immunoprecipitation (ChIP) assays showed that ETS1 is recruited to its binding site on the *ISLR* promoter (Fig 2I). Thus, our data strongly demonstrate that ETS1 activity directly regulates *ISLR* induction in stromal cells.

### Stromal cell-derived Islr is essential for intestinal epithelial regeneration

Next, to investigate the *in vivo* physiological and pathological roles of *Islr*-mediated communication between stromal and epithelial cells in the intestine, we generated *Twist2-Cre*-driven *Islr* cKO mice, in which *Islr* was specifically deleted in stromal cells (**Fig EV2A and B**). The expression level of *Islr* was markedly reduced in the colonic and intestinal tissues of cKO mice (**Fig EV2C and D**). *In situ* hybridization showed that *Islr* was specifically deleted in stromal cells but not in epithelial cells (**Figure EV2E**). Meanwhile, the *Islr* cKO mice were viable and fertile with no apparent gross phenotypes. No differences were found in colon microanatomy, in proliferation and apoptotic patterns within the crypt, or in the number of goblet cells between control and cKO mice (**Fig EV2F-H)**. These results suggest that stromal cell-expressed Islr is dispensable for intestinal homeostasis under physiological condition.

Next, we sought to assess the pathological role of *Islr* in colitis. To this end, we induced experimental colitis in both control and cKO mice by administering DSS in drinking water for 5 days, followed by 3 days of recovery after DSS removal. Body weight did not significantly differ between control and cKO mice during the DSS treatment (Fig 3A). However, after DSS removal, the body weight of cKO mice continue to decrease, while that of control mice appeared to recover towards the starting level (Fig 3A), indicating impaired recovery in the absence of stromal cell-expressed *Islr*. Correspondingly, the inflammatory response was not altered in cKO mice during DSS treatment (Figs 3B **and** C**, and EV2I**), but the regenerative response was impaired after DSS removal (Fig 3B **and** C). In line with this finding, cKO mice exhibited signs of impaired regeneration, with higher clinical scores, shorter colon lengths, and lower percentage of proliferative epithelial cells than the control mice two or three days after DSS removal (Fig 3D-E). Furthermore, an impaired regenerative response upon *Islr* deletion in stromal cells was also found in another mouse model of colitis induced by TNBS (**Figure EV3A-F**). Together, our results indicated that *Islr* plays a key role in promoting epithelial regeneration in colitis.

**Figure 3.**
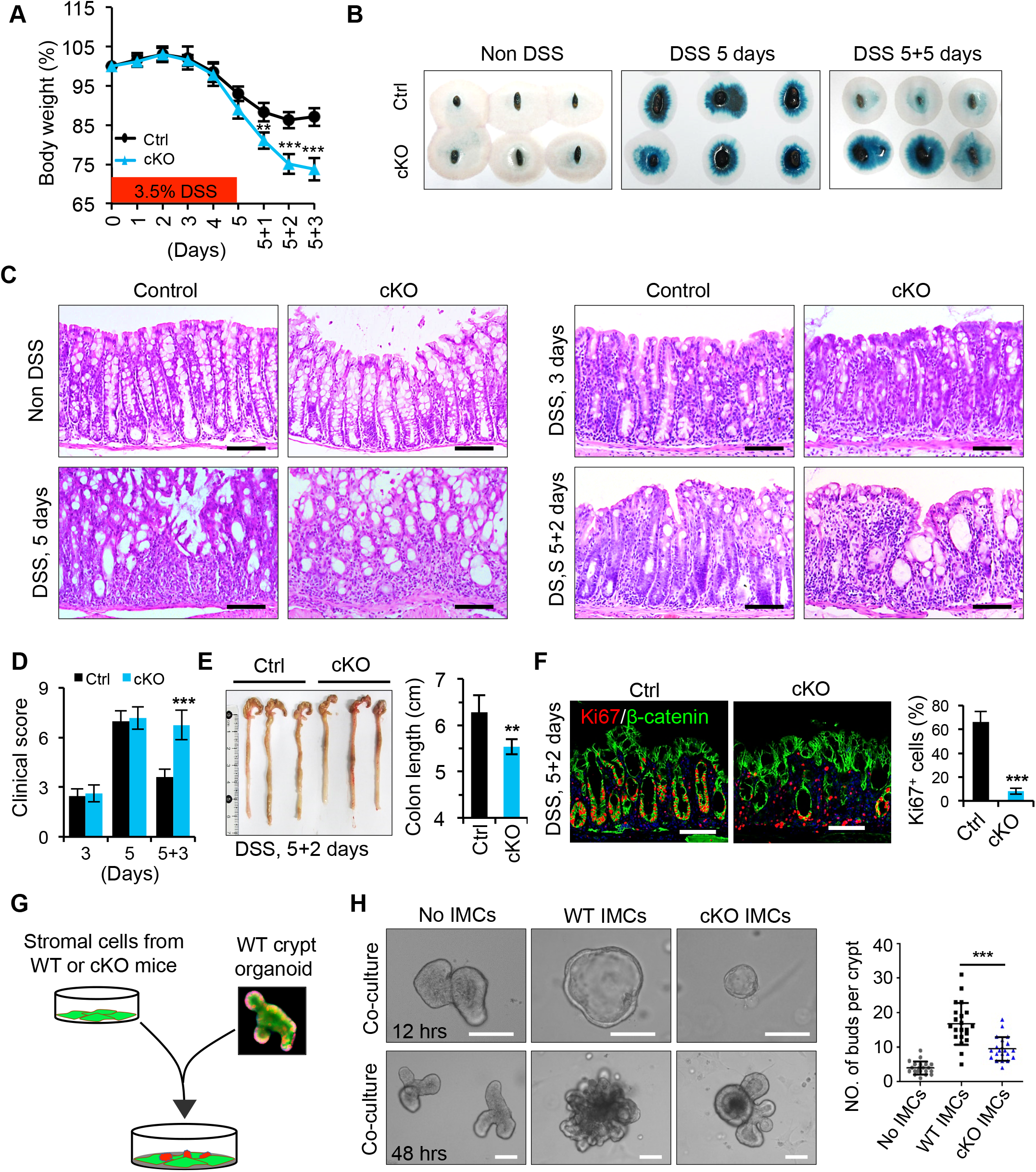
Deletion of *Islr* in stromal cells impaired intestinal regeneration. (A) Quantification of body weight change in control (n = 12) and cKO (n =13) mice after DSS treatment and DSS removal. \*\**P* < 0.01; \*\*\**P* < 0.001. (B) Fecal occult blood test in control and cKO mice at indicated timepoints. (C) Histological images of colonic tissues from control (n=10) and cKO (n=10) mice at indicated timepoints after DSS treatment. Scale bar: 100 μm. (D) Quantification of the clinical scores in control and cKO mice three days after 5-day DSS treatment. \*\**P* < 0.01. (E) Gross images of colons and quantification of colon length from control (n=6) and cKO (n=6) mice two days after 5-day DSS treatment. \*\**P* < 0.01. (F) Double immunofluorescence for Ki67 and β-catenin in colons from control (n=4) and cKO (n=4) mice two days after 5-day DSS treatment. Quantification of percentage of Ki67^+^ epithelial cells. \*\*\**P* < 0.001. (G) The schematic image showing co-culture of WT crypt organoids and intestinal mesenchymal stromal cells (IMCs) from WT and cKO mice. (H) Representative images of WT organoids co-cultured with WT or cKO IMCs for 12 and 48 hours. The WT crypt organoid culture without IMCs treatment (no IMCs) were used as controls. The budding numbers of organoids under indicated conditions after 48-hour co-culture were quantified. \*\*\**P* < 0.001. In A, D-E and H, data are presented as mean ± SD. \*\**P* < 0.01; \*\*\**P* < 0.001 (Student’s t-test).

We also examined the pro-regenerative role of *Islr* in an intestinal organoid model, that partially mimicking intestinal regeneration (Serra et al., 2019). First we isolated intestinal mesenchymal cells (IMCs) from both wild type (WT) and cKO mice. Then, we co-cultured WT crypt organoids with the equal numbers of WT or *Islr*-deficient IMCs. We observed that the organoids grew smaller and produced fewer buddings in *Islr*-deficient IMCs, relative to WT IMCs (Fig 3G **and** H). This supports that *Islr* loss in stromal cells impairs intestinal epithelial regeneration. Taken together, these results demonstrate an essential role for stromal cell-expressed *Islr* in epithelial regeneration especially under inflammation-related pathological conditions.

### Stromal cell-expressed Islr is required for colorectal tumorigenesis

Given the promoting effect of *Islr* on epithelial cell proliferation and its high expression in colorectal cancer, we hypothesized that prolonged Islr expression by stromal cells may exacerbate tumorigenesis. To test this possibility, we examined the *in vivo* role of *Islr* in the AOM-DSS induced colorectal adenocarcinoma model that recapitulates inflammation-driven tumorigenesis (De Robertis, Massi et al., 2011). We found that deletion of *Islr* in stromal cells resulted in marked decreases in both tumor size and tumor number (Fig 4A-D), with concomitant reductions in body weight during the process of tumor induction (Fig 4B). The hyperplastic areas in colonic epithelium are markedly reduced in cKO mice compared to control mice, and cell proliferation levels were significantly lower (Fig 4F-H). In contrast, addition of human recombinant ISLR to the supernatant promoted proliferation of HT29 CRC cells, as compared to non-ISLR control (Fig 4I). Taken together, these results demonstrate that stromal cell-secreted Islr is essential for colorectal tumorigenesis.

**Figure 4.**
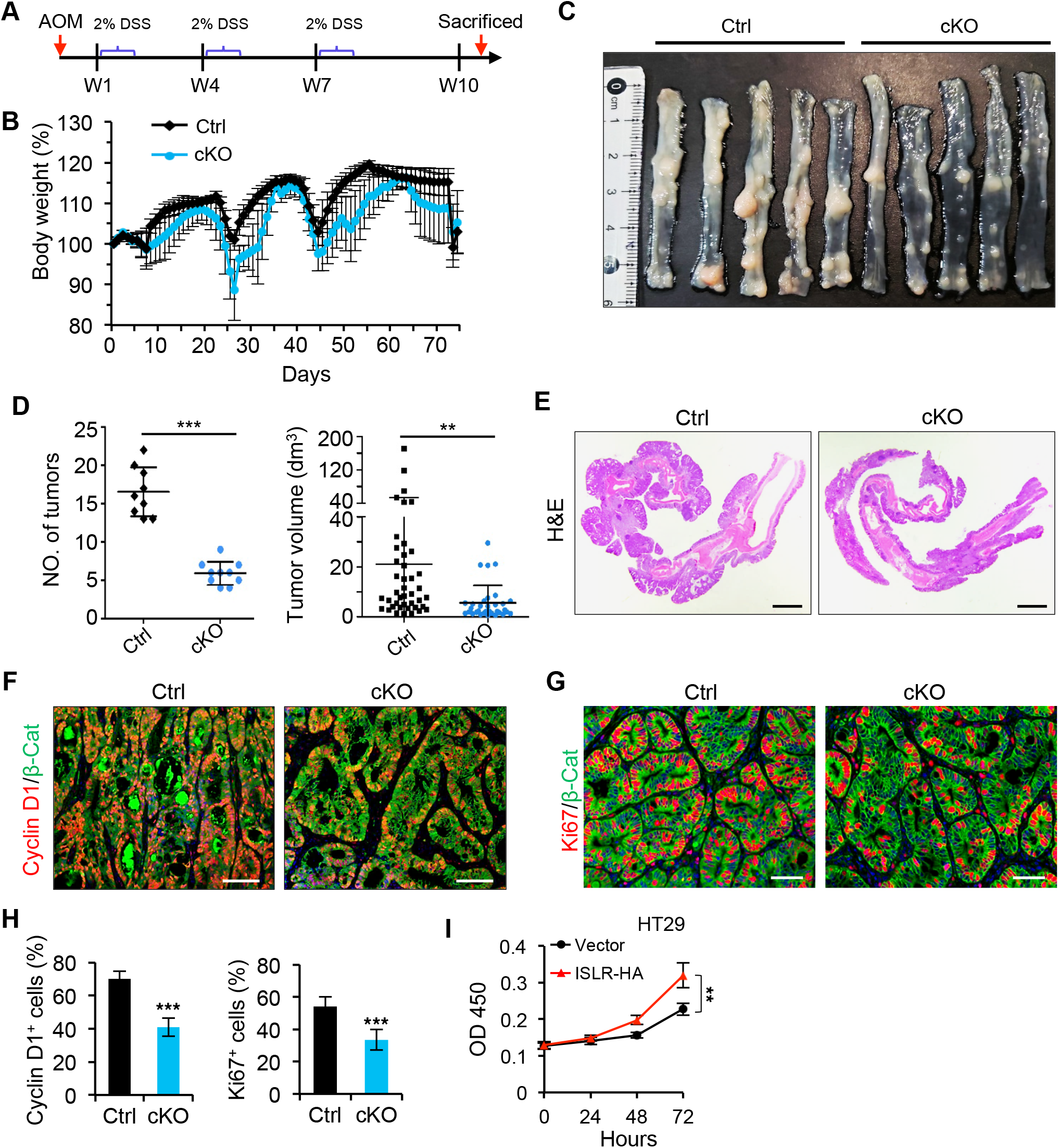
Deletion of *Islr* in stromal cells suppressed tumor growth. (A) Schematics of generating AOM-DSS mouse colon tumor model. (B) Quantification of body weight changes in control and cKO mice during tumor development. (C) Gross images of AOM-DSS tumors in control (n=9) and cKO (n=9) mice. (D) Quantification of tumor number per mouse and tumor volume in control and cKO mice. \*\**P* < 0.01; \*\*\**P* < 0.001. (E) Histological images of colon tumors from control and cKO mice. Scale bar: 2 mm. (F and G) Double immunofluorescence for Cyclin D1 and β-catenin (F), as Ki67 and β-catenin (G) in colon tumors from control and cKO mice. Scale bar: 100 μm. (H) Quantification of percentage of cyclin D1^+^ and Ki67^+^ cells in Panels F and G. \*\*\**P* < 0.001. (I) The growth curve of H29 CRC cells cultured in the supernatant with or without ISLR-HA. The supernatant were collected for H29 CRC cells 24 hours after transfected with pcDNA3.1 empty vector or pcDNA3.1-ISLR-HA vector. \*\**P* < 0.01. In D, H and I, data are presented as mean ± SD. \*\**P* < 0.01; \*\*\**P* < 0.001 (Student’s t-test).

### Stromal Islr dampens epithelial Hippo signal to activate YAP signaling

To gain insight into the molecular mechanism underlying the regulatory effect of Islr on epithelial regeneration and tumor growth, we performed whole genome transcriptome analysis on colonic epithelial cells from control (n=3) and cKO (n=3) mice. We defined the differentially expressed genes (DEGs) as those with *P* < 0.05 and fold change > 2. This approach yielded 996 downregulated genes and 779 upregulated genes in cKO mice (**Fig EV4A**). The Hippo signaling pathway was among the enriched KEGG pathway terms (**Fig EV4B**). We note that YAP1 is a downstream effector of the Hippo pathway and it is essential for promoting intestinal epithelial regeneration and tumorigenesis (Cai et al., 2010, Gregorieff et al., 2015, Serra et al., 2019, Taniguchi et al., 2015, Yui et al., 2018), and that, in particular, *Yap1*-null mice with colitis also exhibit impaired regeneration with no alteration in immune response, thus phenocopying *Islr* cKO mice with colitis (Cai et al., 2010).

Further analysis of genes related to Hippo pathway revealed that many downstream target genes of YAP/TAZ activity, such as *Ctgf* and *Fstl1*, were downregulated, while those upstream inhibitors of the Hippo pathway, such as *Mob1b* and *Dlg3*, were upregulated in cKO mice (Fig 5A). This suggests systematic alteration of Hippo signalling in intestinal epithelium upon stroma-specific *Islr* deletion. These observations are corroborated by qRT-PCR analysis on cKO colon epithelial cells for individual target genes of YAP signaling, including *Ctgf* and *Cyr61* (Fig 5B), as well as by Western blot result for the corresponding proteins (Fig 5C).

**Figure 5.**
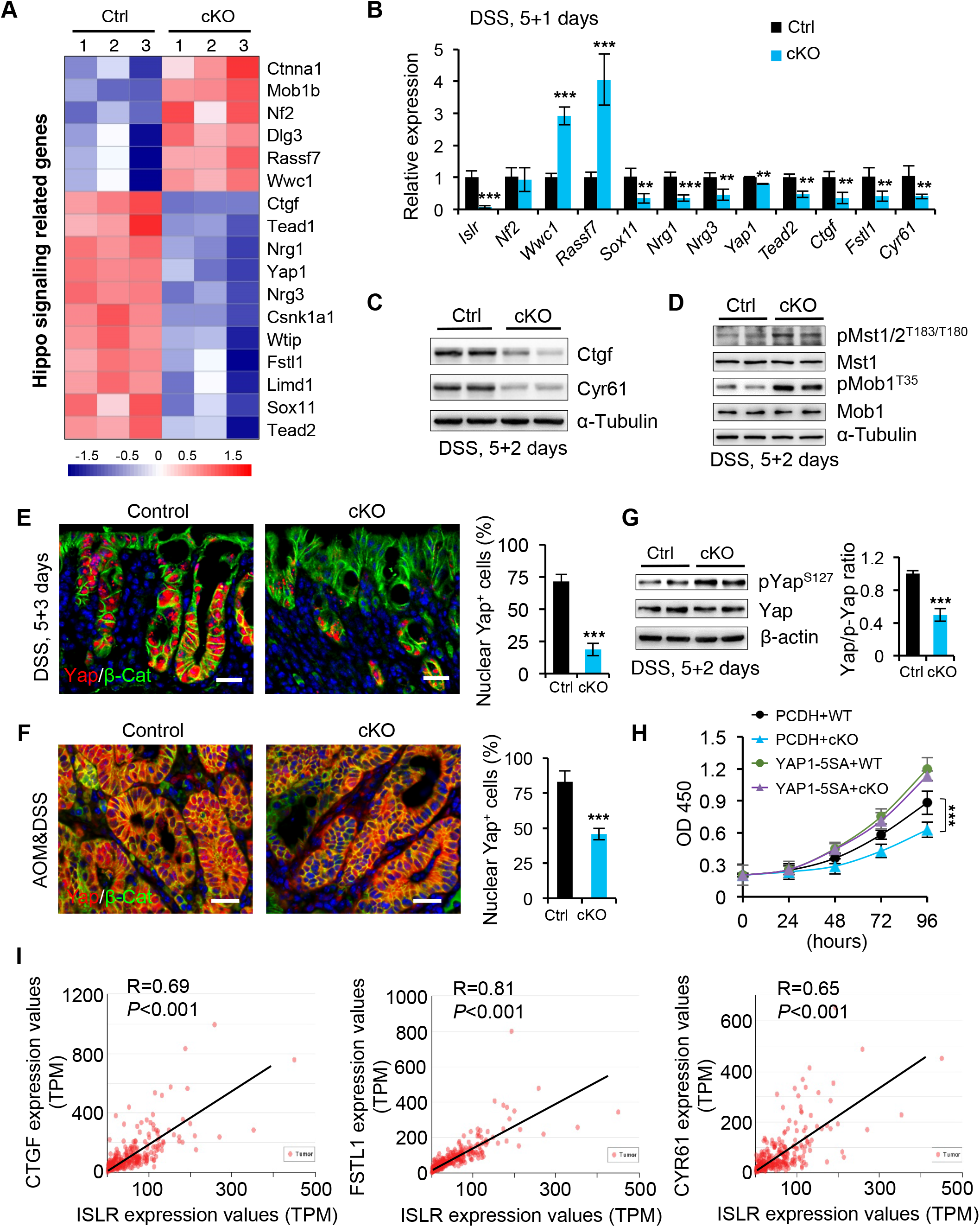
Deletion of *Islr* in stromal cells suppressed epithelial Yap signaling. (A) Heatmap of the altered Hippo related genes in colons from control and cKO mice one day after 5-day DSS treatment. (B) qRT-PCR analysis validates the altered Hippo-related genes in control and cKO mice. n=4. \*\**P* < 0.01; \*\*\**P* < 0.001. (C) Western blotting for Ctgf and Cyr61 in colon tissues from control and cKO mice two days after 5-day DSS treatment. α-Tubulin was used as a loading control. (D) Western blotting for p-Mst1/2, Mst1, p-Mob1 and Mob1 in colon tissues from control and cKO mice two days after 5-day DSS treatment. α-Tubulin was used as a loading control. (E and F) Double immunofluorescence for Yap and β-catenin in the colons from control (n=4) and cKO (n=4) mice three days after DSS removal (E), and in AOM/DSS tumors from control (n=4) and cKO (n=4) mice (F). The percentage of nuclear Yap1^+^ cells versus epithelial cells was quantified. Scale bar: 50 μm. \*\*\**P* < 0.001. (G) Western blotting for Yap and p-Yap in colon tissues from control and cKO mice two days after 5-day DSS treatment. β-actin was used as a loading control. The ratio of Yap::p-Yap were quantified. \*\*\**P* < 0.001. (H) The growth curve of HCT116 colorectal cancer cells cultured in the supernatant from WT or cKO IMCs, concomitantly transfected with PCDH empty vector or PCDH-YAP1-5SA vector. n = 4 technical replicates. \*\*\**P* < 0.001. (I) Pearson correlation analysis of ISLR and CTGF (*P* < 0.001; R = 0.69), ISLR and FSTL1 (*P* < 0.001; R = 0.81), as well as ISLR and CYR61 (*P* < 0.001; R = 0.65) in human colorectal cancer based on TCGA RNA-seq database. In B and E-H, data are presented as mean ± SD. \*\**P* < 0.01; \*\*\**P* < 0.001 (Student’s t-test).

To clarify the regulatory effect of stromal cell-secreted Islr on epithelial Hippo signaling, we examined the activity of Hippo kinases Mst1/2 in the epithelium of DSS-treated mice two days after DSS removal. Our results showed that the phosphorylation of Mst1/2 (pT183/T180 as the marker of kinase activation), as well as that of the substrate Mob1 was markedly increased in the epithelium of cKO mice (Fig 5D). In line with this, immunofluorescence assay for YAP1 revealed a sharp decrease in nuclear YAP1 both in the inflamed mucosa of cKO mice three days after DSS removal (Fig 5E) and in AOM-DSS tumors from cKO mice (Fig 5F). Consistently, phosphorylation levels of Yap1 (pS127) were significantly elevated upon depletion of stromal *Islr* (Fig 5G). Furthermore, we found that the supernatant without recombinant ISLR suppressed cell growth compared to the supernatant with ISLR, while YAP1-5SA-induced YAP activation can rescue the inhibitory effect on cell growth caused by *ISLR* loss (Figs 5H **and EV4C**), suggesting that YAP activity functionally acts as a downstream effector of ISLR.

Given the above findings that ISLR is upregulated in CRC and that stromal depletion of *Islr* results in decreased activity of YAP1, we went further to examine the potential correlation between ISLR and YAP1 activity in clinical samples of human colorectal adenocarcinoma. The transcript levels of *ISLR* are positively correlated with those of YAP1 target genes, *CGTF*, *FSTL1* and *CYR61* (Fig 5I). Taken together, these results demonstrate that stromal ISLR can suppress the Hippo signaling to activate YAP1 in epithelial cells.

### ISLR directly binds epithelial FAT1 to attenuate Hippo signals for YAP activation

To further dissect the mechanism underlying stromal ISLR regulating epithelial Hippo signaling, we asked whether extracellular ISLR secreted by stromal cells could suppress the Hippo signaling in epithelial cells *in vitro*. To this end, we first overexpressed HA-tagged human ISLR (hISLR-HA) in HEK293FT cells and confirmed that this results in an increase in secreted ISLR in the supernatant (Fig 6A). We then treated HEK293FT cells using the supernatant with or without hISLR-HA and performed RNA transcriptome analysis. Hippo signaling pathway was also among the enriched KEGG pathway terms (**Fig EV5A**). Gene set enrichment assay (GSEA) showed that a set of direct YAZ/TAZ/TEAD target genes was significantly enriched among the DEGs upon hISLR-HA treatment (Fig 6B **and** C), suggesting that extracellular hISLR triggers the activation of YAP signaling. In line with this, the levels of pMST1/2, pMOB1 and pYAP1 were reduced in a dose-dependent manner upon extracellular hISLR-HA (Fig 6D).

**Figure 6.**
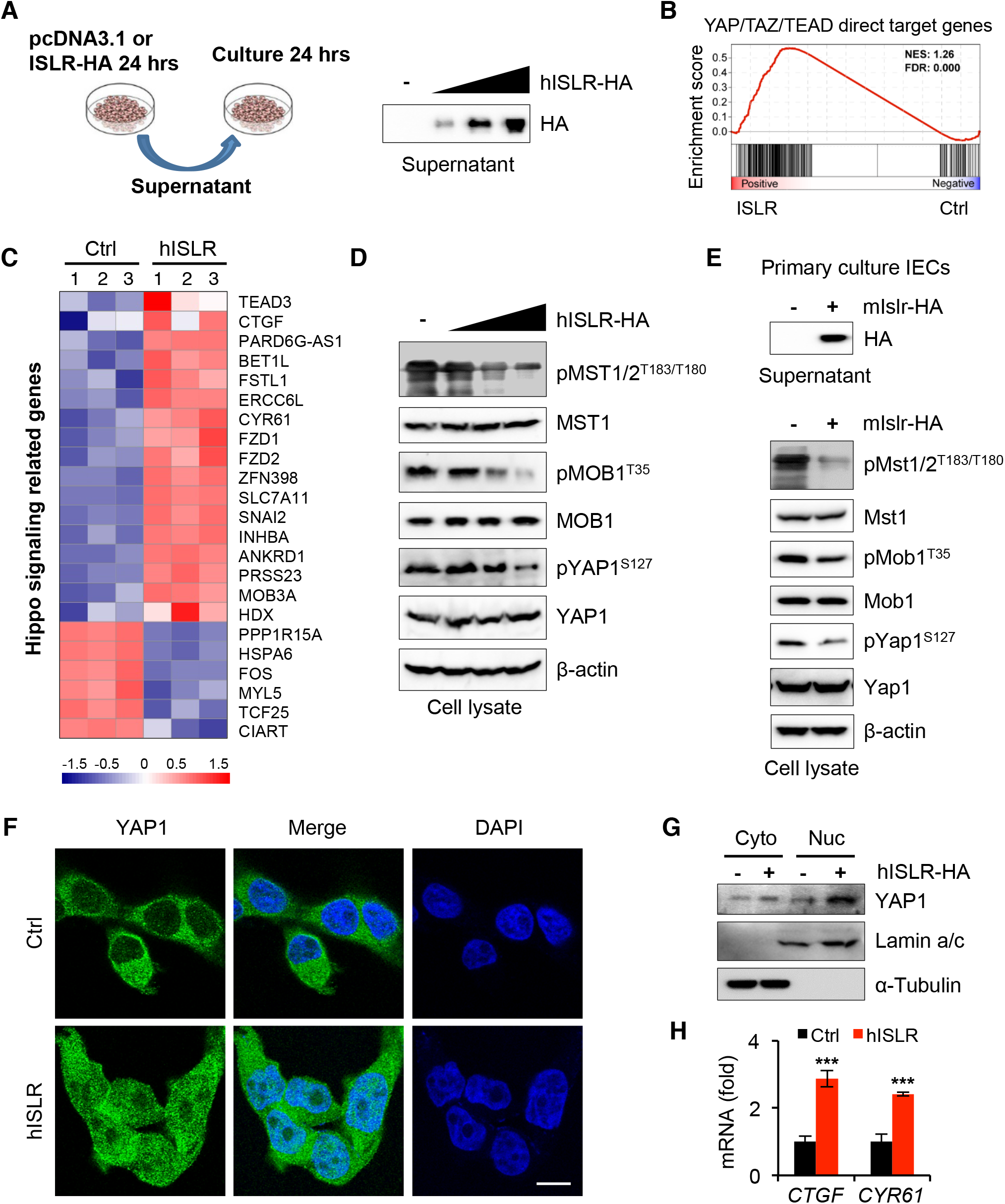
Extracellular Islr attenuates epithelial Hippo signaling to activate Yap signaling *in vitro*. (A) Schematics of culture epithelial cells with supernatant from HEK293FT cells transfected with pcDNA3.1 empty vector or pcDNA3.1-hISLR-HA plasmids. Western blotting for HA in the supernatant from HEK293FT cells transfected with the indicated plasmids. (B) Gene set enrichment for YAP/TAZ/TEAD direct target genes in hISLR-HA overexpressing transcriptome profiles. The transcriptome analysis was performed in HEK293FT cells transfected with the indicated plasmids. (C) Heatmap showing the altered Hippo related genes in the hISLR-HA overexpressing transcriptome profiles. (D) Western blotting for pMST1/2, MST1, pMOB1 and MOB1, pYAP1 and YAP1 in lysates from HEK293FT cells cultured in the supernatants with or without hISLR-HA. β-actin was used as a loading control. (E) Western blotting for HA in the supernatant from primary cultured intestinal epithelial cells (IECs) transfected with pcDNA3.1 or pcDNA3.1-hISLR-HA plasmids, and western blotting for pMST1/2, MST1, pMOB1 and MOB1, pYAP1 and YAP1 in lysates from primary IECs cultured in the supernatants with or without hISLR-HA. β-actin was used as a loading control. (F) Immunofluorescence for YAP1 in primary intestinal epithelial cells cells cultured in the supernatant with or without hISLR-HA. Scale bar: 10 μm. (G) Western blotting for YAP1 in cytoplasmic or nuclear proteins from HEK293FT cells cultured in the supernatant with or without hISLR-HA. Lamin a/c, a positive control for nuclear proteins. α-Tubulin, a positive control for cytoplasmic proteins. (H) qRT-PCR for *CTGF* and *CYR61* in HEK293FT cells cultured in the supernatant with or without hISLR-HA. \*\*\**P* < 0.001. In G, data are presented as mean ± SD. \*\*\**P* < 0.001 (Student’s t-test).

Importantly, reduction of pMST1/2, pMOB1 and pYAP1 upon extracellular hISLR-HA treatment was further confirmed in primary culture of intestinal epithelial cells (Fig 6E), in NCM460 intestinal epithelial cells, and in HCT116 CRC cells (**Fig EV5B-F**). Consistent with the above findings, treatment with the extracellular hISLR-HA markedly increased the nuclear translocation of YAP1 in HEK293FT and NCM460 epithelial cells, evidenced by fluorescent staining and Western blot (Fig 6F **and** G). Meanwhile, expression of YAP1 target genes *CTGF* and *CYR61* were also significantly increased in response to the extracellular hISLR-HA treatment (Fig 6H). Together, these results clearly demonstrate that extracellular hISLR directly attenuates Hippo signaling in target cells, and consequently triggering YAP activation.

To probe which specific molecule(s) on the surface of target cells mediate the attenuating effect of secreted ISLR on Hippo signaling, we performed co-immunoprecipitation (Co-IP) using specific anti-ISLR antibody, followed by shortgun Mass Spectrometry to identify ISLR binding partners (Maccarrone, Bonfiglio et al., 2017). Our results identified a group of putative ISLR-binding partners, among which MAGT1, FAT1, FCGR3B, ATP1B3 rank the highest (Fig 7A **and** B). Noting that FAT1 is a transmembrane protein that was previously reported as an upstream regulator of Hippo pathway and could recruit and activate MST1/2 kinases (Ahmed et al., 2015, Martin et al., 2018), we hypothesized that the secreted ISLR directly interacts with FAT1 to interfere with the Hippo signaling inside the target cells. To test this hypothesis, we carried out Co-IP analysis, which confirmed that ISLR can directly bind to FAT1 (Fig 7C). Further domain mapping identified that the extracellular region of FAT1 corresponding its Cadherin domain 13-15 (amino acids 1457-1765) is essential for binding ISLR (Figs 7D **and EV5G**). Moreover, the Ig-like domain of ISLR (amino acids 230 to 350) is required for binding FAT1 (Fig 7E), as well as suppression of Hippo signaling (Fig 7F). Finally, we examined whether ISLR binding could affect the association between FAT1 and MST1. Our Co-IP result showed that ISLR treatment decreased the association of MST1 with FAT1 in a dose-dependent manner (Fig 7G). Taken together, these results indicate that stromal cell-secreted ISLR binds to FAT1 on the plasma membrane of epithelial cells to impair the recruitment and activation of MST1/2 kinases inside the cells, thus attenuating Hippo signaling in the epithelial cells.

**Figure 7.**
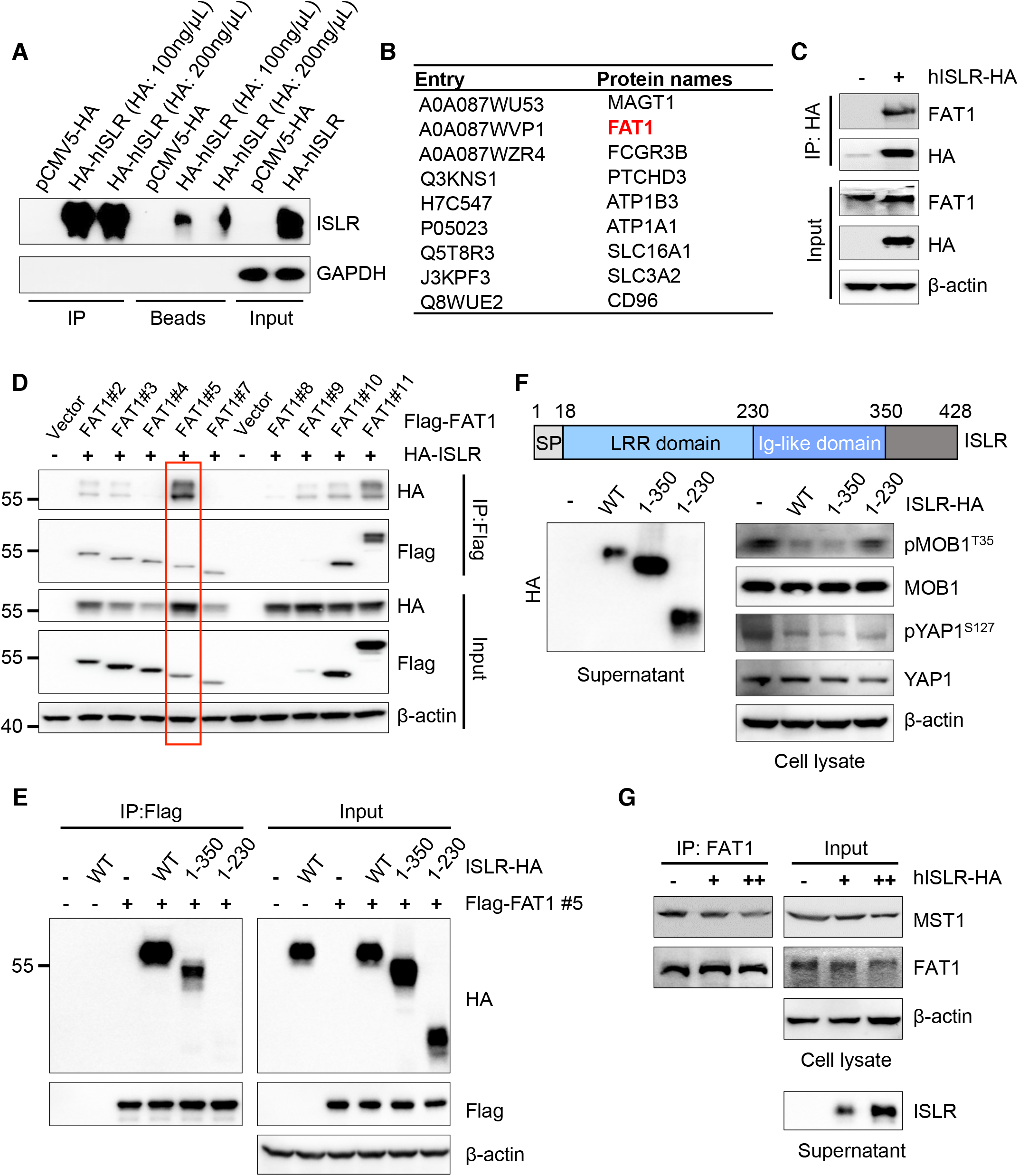
ISLR directly binds to a transmembrane protein FAT1, decreasing the association of MST1 with FAT1. (A) Western blotting with ISLR antibody in lysates of HEK293FT cells transfected with pCMV5-HA and pCMV-hISLR-HA plamids, and in elution collected from beads, after immunoprecipitation with HA antibody at concentrations of 100 or 200 ng/μL. GAPDH was used as a control for the input. (B) A list of ISLR binding transmembrane protein partners were identified by Co-IP-coupled shortgun Mass Spectrometry. (C) Co-IP analysis of ISLR and endogenous FAT1 in HEK293FT cells transfected with the ISLR-HA plasmids. Expression vectors encoding hISLR-HA was transfected into HEK293FT cells. β-actin was used as a loading control. (D) Domain mapping by Co-IP to identify the specific region of FAT1 (amino acids 1457-1765) that interacts with ISLR. The information of FAT1 segments was shown in supplementary Figure 5G. (E) Domain mapping to identify the specific region of ISLR (amino acids 230-350) responsible for binding FAT1. (F) Schematic representation of the domain organization of ISLR (Upper). SP, signaling peptide; LRR, leucine-rich repeats. Protein levels of pMOB1, MOB1, pYAP1 and YAP1 in HEK293FT cells treated with supernatants containing different ISLR segments were detected with the indicated antibodies (lower). β-actin was used as a loading control. The supernatants of different ISLR segments were detected with HA antibody, shown in the left panel. (G) Co-IP analysis of the interactions between endogenous FAT1 and MST1 in HEK293FT cells treated with or without ISLR supernatant.

## Discussion

Here we demonstrate that stromal cell-secreted ISLR promotes intestinal epithelial regeneration by attenuating Hippo signaling to activate Yap signaling in epithelial cells (Fig 8). Consistent with this conclusion, numerous studies showed an essential role for YAP/TAZ in intestinal regeneration upon inflammation or after irradiation (Cai et al., 2010, Gregorieff et al., 2015, Serra et al., 2019, Taniguchi et al., 2015, Yui et al., 2018). Supporting the function of ISLR in regulating Hippo signaling, Yap1 deficiency in the intestine of mice with colitis impairs the regenerative response but does not influence the inflammatory response (Cai et al., 2010). Thus, it results in a phenotype that resembles that of the *Islr* cKO mice. Given the central importance of YAP/TAZ in tissue repair, it is conceivable that a sophisticated upstream regulatory network is necessary for fine-tuning YAP/TAZ activity. In this regard, recent reports have indicated that YAP/TAZ signaling can be activated in colitis by the IL-6/GP130/Src as well as extracellular matrix (ECM)/FAK/Src cascades (Taniguchi et al., 2015, Yui et al., 2018), independent of the classic Hippo pathway. Our current work shows how epithelial Hippo signaling is directly regulated by secreted ligand of stromal origin upon inflammatory stimuli. Therefore, we now define an inter-cellular regulatory machinery for the classic Hippo-YAP pathway in the epithelial cells that is governed by signals from stromal cells, and provides a parallel mechanism accounting for YAP-associated cell-cell communications. In addition to highlighting the importance of stromal cells in controlling epithelial YAP activity under pathological conditions, our study also identifies novel ETS1-ISLR−FAT1-Hippo signaling axis that links inflammation-associated stromal signals with epithelial regeneration.

**Figure 8.**
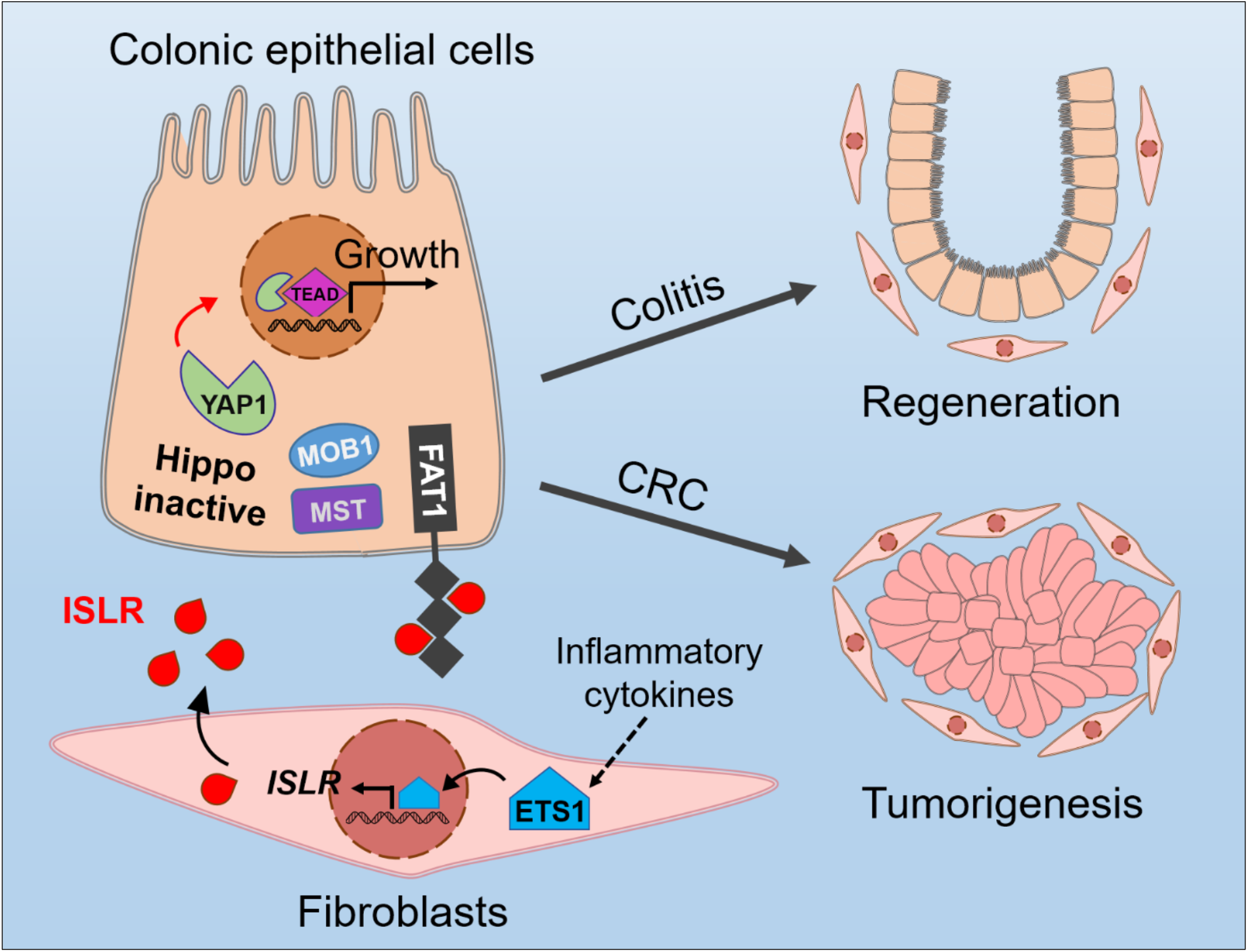
The working model of stromal cell-secreted Islr regulating intestinal epithelial Hippo signaling.

Previous reports have shown that FAT1 is a transmembrane protein acting as an upstream activator of Hippo kinase (Li et al., 2018, Martin et al., 2018). In this work, we show that ISLR can directly bind to the extracellular region of FAT1 to attenuate the Hippo signals inside of epithelial cells, thus activating YAP. Although the exact mechanism of how FAT1 relays ISLR signal to MST1/2 kinases is not clear, it is most likely that ISLR binding triggers an outside-in conformational change of FAT1, thus impairing its recruitment and activation of MST1/2. Thus, it appears that ISLR functions as an extracellular ligand of FAT1, and that FAT1 acts as a cell surface receptor of the Hippo pathway. Previously, GPCRs and CD44 have been proposed as potential cell surface receptors of Hippo signaling that mediate extracellular signals (Xu, Stamenkovic et al., 2010, Yu, Zhao et al., 2012). As of now, the upstream regulatory mechanisms of the Hippo pathway, especially the ones mediated by extracellular signals in a context-dependent manner remain incompletely understood. In this regard, our current work identifies a molecular mechanism for the regulation of Hippo signaling across two major cell types in the intestine. Considering the marked upregulation of stromal ETS1 and ISLR, and the significant alteration of epithelial Hippo signals and YAP activity in the context of colitis and cancer, the ETS1-ISLR−FAT1-Hippo signaling axis connecting stroma to epithelium may represent a major molecular mechanism that is responsive to stress conditions.

Finally, our observations that deletion of *Islr* in stromal cells suppressed tumor growth and that *ISLR* levels are positively correlated with the poor survival and lymph node metastasis, strongly support the idea that ISLR acts as an oncogenic factor during CRC development. Other pro-tumorigenic factors secreted by tumor-associated stromal cells, including IL6, VEGF and MMPs, have been shown to promote tumorigenesis (Bussard et al., 2016). Intriguingly, our results highlighted ISLR as a stroma-secreted oncogenic factor that can enhance YAP activity in CRC. Our finding of the inter-cellular connection between ISLR and YAP/TAZ explains the molecular basis for TAP/TAZ hyperactivation commonly observed in many types of human cancers (Overholtzer, Zhang et al., 2006). In line with its oncogenic activity, ISLR was found in this work to directly bind and suppress the tumor suppressor activity of FAT1, which is frequently mutated in various types of cancers (Li et al., 2018, Martin et al., 2018). Consistently, we demonstrate that ETS1, which is an oncogenic transcription factor in most cancer types (Dittmer, 2015), is a direct upstream regulator of *ISLR* and it increases its expression in response to stress signals. In support of this notion, both ISLR and ETS1 are upregulated in tumor-associated stromal cells(Wernert, Gilles et al., 1994), and their expression levels are correlated with tumor progression (Dittmer, 2015). The ETS1-induced ISLR expression can at least partially account for the oncogenic activity of ETS1. Thus, we posit that ISLR is an important pro-tumorigenic factor in CRC.

In summary, we discovered a new molecular mechanism underlying communication between stromal cells and epithelial cells during intestinal tissue regeneration and tumorigenesis. This mechanism primarily features ISLR (a stromal ligand)-FAT1 (an epithelial receptor) interaction that attenuates Hippo signaling to allow for Yap activation in the epithelium. While this type of stroma-epithelium signaling is required for regeneration, its dysregulation exacerbates colorectal cancer– a finding that offers basis for a new therapeutic strategy.

## Materials and Methods

### Subjects

Colonoscopic biopsies were obtained from inflamed and unaffected sites in the colons of 10 UC and 10 CD patients as well as from the normal colonic mucosa in 10 healthy individuals. All patients were from Department of Gastroenterology, the Shanghai Tenth People’s Hospital of Tongji University (Shanghai, China). Each subject provided written informed consent. The diagnosis of UC and CD was based on clinical, radiological, endoscopic and pathological examinations.

### Ethics

All mouse experimental procedures and protocols were evaluated and authorized by the Beijing Laboratory Animal Management, and were strictly performed in accordance with the guidelines of the Institutional Animal Care and Use Committee of China Agricultural University (approval number: SKLAB-2015-04-03).

### Mouse strains

The *Islr* floxed mouse has been described previously (Zhang et al., 2018). The *Twist2-Cre* mice were obtained from the Jackson Laboratory (stock number: 008712).

### RNAscope in situ hybridization assay

An RNAscope *in situ* hybridization assay was performed as described previously (Anderson, Zhang et al., 2016). The sample sections were processed according to the manufacturer’s instructions using an RNAscope Red kit (ACDBio). Briefly, 5 µm thick sections were deparaffinized, and treated with hydrogen peroxide solution for 10 min at room temperature (RT). Target retrieval was performed for 15 min at 100°C, and then the slides were treated with protease for 15 min at 40°C. Probes were then hybridized for 2 hours at 40°C, followed by RNAscope amplification and then Fast Red chromogenic detection RNAscope. Then the slides were counterstained in hematoxylin. After dehydration, the slides were mounted with EcoMount (EM897L; Biocare). The following RNAscope probes were used in this study: Hs-ISLR (cat. #455481; ACDBio), Mm-Islr-O1 (cat. #453321; ACDBio), Mm-Ppib (cat. #313911; ACDBio), DapB (cat. #310043; ACDBio).

### Co-IP assay

Lysates were prepared by incubating cells on ice with cold TBS-Nonidet P-40 lysis buffer (20 mM Tris, pH 7.4, 150 mM NaCl, and 1% Nonidet P-40) in the presence of a protease and phosphatase inhibitor cocktail (Roche) for 30 min at 4°C. Then, the cells were scraped and centrifuged at 12,000 rpm for 10 min at 4°C. For immunoprecipitation, proteins (approximately 0.5 mg) were incubated with control or specific antibodies (3–5 μg) for 12 hours at 4°C under constant rotation. Then, 30 μL of pretreated protein G magnetic beads (Santa Cruz) were added, and the mixture was incubated overnight at 4°C. The beads were washed three times with lysis buffer and then collected by centrifuging at 4°C. The proteins were eluted by boiling for 10 min with 2× SDS-PAGE loading buffer. The immune complexes were analyzed by SDS-PAGE using appropriate antibodies.

### DSS treatment

To induce acute colitis with DSS, mice were treated with 3.5% w/v DSS with molecular weight of 36,000–50,000 (MP Biochemicals) in drinking water for 5 days; and then the DSS was withdrawn for another 3 days. The severity of colitis was scored daily on the basis of standard parameters, including body weight and the presence of diarrhea, and/or bloody stools. The mice were sacrificed at different time points to obtain colon samples. The clinical score was measured as previously reported with modifications (Tian, Xu et al., 2019), and evaluated by an independent investigator who was blinded to the experiment. The parameters of clinical score are included in Appendix table 1.

### TNBS treatment

TNBS was used to induce colitis as described previously (Tian et al., 2019). To induce colitis with TNBS, the mice were weighed and anesthetized with isoflurane-mixed gas on day 1. Then, a 1.5 x 1.5 cm area of skin was shaved on the back of each mouse between the shoulders, and 150 μL of 1% (w/v) TNBS (Sigma, cat. no. p2297) presensitization solution was applied to the shaved skin. On day 8, after weighing and anesthetizing each mouse, we slowly injected 125 μL of 2.5% (w/v) TNBS solution into the lumen of the colon with 1 mL syringe and a 3.5 F catheter, and removed the catheter and positioned the mouse head-down for 5 minutes. The severity of colitis was scored daily on the basis of standard parameters including body weight and the presence of diarrhea and/or bloody stools. The mice were sacrificed at the indicated timepoints to obtain colon samples.

### AOM and DSS treatment

An AOM-DSS mouse model was generated as described previously with modifications (Tian, Ma et al., 2017). Briefly, eight-week-old control and *Islr* cKO mice were intraperitoneally injected with AOM at a concentration of 10 mg/kg body weight (Sigma-Aldrich). Five days after AOM injection, the mice were treated with a so-called DSS cycle comprising two steps: the mice were first given 2.5% (w/v) DSS (molecular weight 36,000–50,000, MP Biomedicals) for 7 days and then given normal water for 14 days. The mice were subjected to a total of 3 DSS cycles. After the treatments, the mice were sacrificed, distal colon tissues were collected, and tumor numbers and volumes were evaluated.

### qRT-PCR analysis

Total RNA was isolated from sorted cells of colonic tissues, cell lines and mouse colonic tissues using TRIzol reagent (Life Technologies) according to the manufacturer’s instructions. To detect mRNA levels, reverse transcription was carried out using oligo (dT) primers. qRT-PCR was performed using LightCycler 480 SYBR Green I Master Mix on a LightCycler 480 Real-Time PCR System (Roche, Mannheim, Germany). Relative expression was calculated based on the 2^−ΔΔCt^ method and GAPDH was used as the internal control. The primers for qRT-PCR analysis are included in Appendix table 2.

### Flow cytometry and cell sorting

Colonic epithelial cells were collected following vigorous shaking after incubation in 1× DPBS containing 5 mM EDTA and 0.2 mM DTT for 30 min at 37°C on a rotating platform. The remaining tissue was incubated in digestion solution containing 2% FBS, 0.25 mg/mL collagenase XI (Sigma) and 0.1 mg/mL DNase I (Sigma) for 1 hour at 37°C. Mesenchymal cells were collected by centrifugation following vigorous shaking. Then, the colonic epithelial cells and mesenchymal fractions were pipetted with prewarmed dispase (2.5 U/mL) and DNase I for 5 min to generate single-cell suspensions. The single-cell suspensions from the colon tissues were stained for 30 min on ice using a mix of antibodies (from eBioscience: LIVE/DEAD-BV421, CD45-APC, Epcam-FITC, and CD90-PE). Viable cells were gated by forward and side scatter; Epcam^+^, Epcam^−^CD45^+^, CD45^−^Epcam^−^CD90^−^, and CD45^−^Epcam^−^CD90^+^ cell subpopulations were gated; and 1 × 10^5^ to 1 × 10^6^ cells were sorted using a BD FACS Aria III sorter (BD Biosciences) for further RNA isolation.

### Isolation and culture of primary IMCs

IMCs were isolated as described previously with modifications (Khalil, Nie et al., 2013). Briefly, colonic tissues were removed and placed in cold HBSS to remove debris. Each colon was opened longitudinally and cut into 0.5 cm pieces, which were then incubated in HBSS containing 5 mM EDTA for 30 min at 37°C on a rotating platform to remove primary intestinal epithelial cells. After isolating the epithelial cells by vigorous shaking, the remaining tissue was rinsed several times with cold HBSS and incubated in digestion solution containing 2% FBS, 0.25 mg/mL collagenase XI (Sigma), 0.08 U/mL dispase II and 0.1 mg/mL DNase 1 (Sigma) for 1 hour at 37°C on a rotating platform. After vigorous shaking, single mesenchymal cells were pelleted at 500 × *g* in a tabletop centrifuge for 5 min. The supernatant was discarded, and each pellet was resuspended with modified DMEM (Gibco) containing 10% FBS, 100 U/mL penicillin, 100 μg/mL streptomycin (Sigma), 1 × L-glutamine and 1 × nonessential amino acids (NEAAs). Then, the cells from each mouse colon were seeded in a 6 cm dish and cultured in a 5% CO_2_ incubator at 37°C. After 2.5 hours, the floating debris and nonadherent cells were washed off with PBS. The adherent cells were IMCs consisting of fibroblasts and myofibroblasts. The IMCs were passaged one week post seeding with a 1:3 ratio. After 3 passages, the supernatant was collected from each dish for follow-up experiments.

### Crypt organoids and IMCs culture

Crypts were isolated from small intestine samples by incubation for 10 min at 4°C in HBSS containing 5 mM EDTA as described previously (Sato, Vries et al., 2009). Briefly, the isolated crypts were counted and pelleted. A total of 100 isolated crypts from C57 BL/B6 mice were mixed with 30 µL of Matrigel (BD Bioscience) and plated in 48-well plates. After Matrigel polymerization, 300 µL of crypt culture medium (advanced DMEM/F12 [Gibco], 100 U/mL penicillin, 100 μg/mL streptomycin [Sigma], B27 supplement [Invitrogen], N2 supplement [Invitrogen], 2 mM GlutaMAX [Invitrogen], 1 mM N-acetyl cysteine [Sigma], 50 ng/mL mouse EGF [PeproTech], 100 ng/mL mouse Noggin [PeproTech] and 10% human R-spondin-1 [PeproTech]) was added to each well. The medium was changed every two days, and the organoids were passaged every four days.

Briefly, isolated crypts were counted and pelleted. A total of 100 isolated crypts from C57 BL/B6 mice were mixed with 30 μL of matrigel (BD Bioscience) and plated in 48-well plates. After matrigel polymerization, 300 μL of crypt culture medium [advanced DMEM/F12 (Gibco), 100U/mL penicillin, 100 μg/mL streptomycin (Sigma), B27 supplement (Invitrogen), N2 supplement (Invitrogen), 2mM Glutamax (Invitrogen), 1 mM N-acetyl cysteine (Sigma), 50 ng/mL mouse EGF (Peprotech), 100 ng/mL mouse Noggin (Peprotech) and 10% human R-spondin-1 (Peprotech)] was added to each well. Medium was changed every two days and organoids were passaged every four days.

For coculture of crypt organoids and IMCs, a total of 50 crypts were mixed with 30 μL of Matrigel and plated in the center of a 48-well plate. After Matrigel polymerization, 300 μL of crypt culture medium containing 1,000 IMCs was added to each well. Crypt organoid growth was observed daily.

### Cell lines, culture conditions and transfections

HEK293FT and HT29 cell lines were purchased from the American Type Culture Collection (ATCC) (Manassas, VA) and maintained in DMEM supplemented with 10% FBS. HT29 cell line was purchased from ATCC (Manassas, VA) and cultured in McCoy’s5a supplemented with 15% FBS. HCT116 and NCM460 cell lines were purchased from ATCC (Manassas, VA) and cultured in RPMI 1640 supplemented with 10% FBS, or in IMDM supplemented with 10% FBS. All cell lines were tested and confirmed to be free of mycoplasma infection. The cells were transfected using Lipofectamine 2000 reagent (Invitrogen) with 2 µg of vector or negative control vector according to the manufacturer’s protocol.

### Histology, immunofluorescence and immunohistochemistry

For histological analysis, the colonic tissues were rinsed with 1x DPBS, fixed in 10% formalin, paraffin-embedded and sectioned at 5 μm. The sections were deparaffinized with xylene followed by treatment with serial dilutions of ethanol. The sections were stained with hematoxylin (Sigma) for 6 min and then washed in running water for 5 min. The sections were stained with eosin (Sigma) for 10 seconds, dehydrated with serial dilutions of ethanol and then mounted with coverslips in neutral gum mounting medium.

For immunostaining, antigen retrieval was performed by heating the slides in 0.01 M citrate buffer (pH 6.0) in a microwave. The sections were rinsed three times with ddH_2_O after natural cooling, immersed in 3% H_2_O_2_ for 10 min or PBS-T (PBS, 1% Triton X-100) for 20 min, washed twice with PBS and then blocked for 1 hour at RT with blocking solution (10% normal goat serum in TBS-T). The sections were incubated with primary antibodies overnight at 4°C. Then, the slides were washed three times with PBS and incubated for 1 hour at RT with Alexa Fluor 488 and 594 goat anti-mouse or anti-rabbit IgG (H+L). The slides were then washed 3 times with PBS and stained with DAPI for 5 min, and the sections were covered with anti-quenching reagent. The following antibodies were used: anti-Ki67 (1:500, Thermo Fisher), anti-ISLR (1:200, Sigma), anti-Ets1 (1:800, Cell Signaling Technology), anti-Sox9 (1:1,000, Cell Signaling Technology), anti-CD45 (1:200, Santa Cruz), anti-F4/80 (1:200, Santa Cruz), anti-β-catenin (1:1,000, Sigma), anti-Cleaved caspase3 (1:1,000, Cell Signaling Technology), anti-Cyclin D1 (1:1,000, Cell Signaling Technology), anti-Yap (1:500, Cell Signaling Technology), anti-Mucin2 (1:100, Santa Cruz), and anti-pSTAT3 (1:1,000, Cell Signaling Technology).

### Nucleoprotein extraction

Nucleoprotein was isolated from cells using a nucleoprotein extraction kit (AR0106, Bosterbio, USA) according to the manufacturer’s instructions. Briefly, HEK293T or NCM460 cells were washed once with cold PBS. The cells were pelleted at 600 × *g* for 5 min, resuspended in reagent A and incubated on ice for 10 min to lyse the cell membrane. Intact nuclei were pelleted at 16,000 × *g* for 5 min, and soluble cytoplasmic proteins were removed. The nuclear pellet was resuspended in nucleoprotein extraction reagent and incubated on ice for 40 min. Chromatin was pelleted at 20,000 × *g* for 5 min, and the soluble nucleoproteins were saved. The protein concentration was determined with a BCA kit (Beyotime). Lamin a/c or Histone H3 was used as a nuclear internal control.

### Western blotting

Cell lysates were subjected to western blotting according to standard procedures. Fresh tissues were homogenized using RIPA buffer (Beyotime Biotechnology, Shanghai, China) in the presence of protease and phosphatase inhibitor cocktails (Roche), followed by treatment with Homogenizer workcenter (T10 basic, IKA). Proteins were measured by BCA protein assay kit (Beyotime) and denatured, then 30 μg of total protein was separated on a 6-12% SDS-PAGE gel, transferred to PVDF membranes (GE Healthcare). The PVDF membranes were blocked with 5% nonfat dry milk for 1 hour at RT and incubated with primary antibodies overnight at 4°C. Results were collected by chemiluminescence imaging system (Sagecreation, Beijing). The relative protein bands intensity were quantified using ImageJ software (U.S. National Institutes of Health, Bethesda, MD, USA). Antibodies were applied as following: anti-GAPDH (Sigma), anti-α-Tubulin (Sigma), anti-β-actin (Sigma), anti-ISLR (Sigma), anti-Histone H3 (Santa Cruz), anti-E-cadherin (Abcam), anti-Lamin a/c (Cell Signaling), anti-HA (Santa Cruz), anti-ETS1 (Cell Signaling), anti-FAT1(Santa Cruz), anti-CYR61 (Santa Cruz), anti-CTGF(Santa Cruz), anti-MOB1(Cell Signaling), anti-pMOB1^T35^ (Cell Saignaling), anti-YAP (Cell Signaling), anti-pYAP1^S127^ (Cell Signaling), anti-pMST1/2^T183/T180^ (Cell Signaling) and anti-MST1(Cell Signaling).

### Plasmid construction

Constructs of full-length or truncated human ISLR and mouse Islr were cloned into a pMlink vector and a pcDNA3.1 vector with a C-terminal 3 × HA tag. FAT1 segments were cloned into a pcDNA3.1 vector with an N-terminal Flag tag and ISLR segments were cloned into a pcDNA3.1 vector with a C-terminal 3 × HA tag. A full-length mouse Ets1 construct was cloned into a modified pCMV5 vector with a C-terminal 3 × HA tag. YAP(5SA)(S61/S109/S127/S164/S381A) plasmids were constructed in a pCDH1-MCS-coGFP vector. For luciferase assays, a construct including 2 Kb Islr promoter sequence was cloned into a pGL3-Basic vector. All mutants were generated through site-directed mutagenesis. All constructs were verified by performing DNA sequencing.

### Microarray analysis

Total RNA was isolated from the colons of three littermate controls and three cKO mice using TRIzol reagent (LifeTechnologies) according to the manufacturer’s instructions. RNA was purified with an RNAse-free DNAse kit (Qiagen) and then submitted to CapitalBio Technology, where the samples were labeled and hybridized to Agilent Mouse Gene arrays. The array data were analyzed for data summarization, normalization and quality control using the GeneSpring software V13 (Agilent). The q value, a measure of the false discovery rate (FDR), was computed using the Significance Analysis of Microarrays software for each probe set by running an unpaired *t*-test. The data were Log2-transformed and the median was centred by genes using the Adjust Data function of Cluster 3.0 software; the data were then further analysed by hierarchical clustering with average linkages. Finally, tree visualization was performed using Java TreeView (Stanford University School of Medicine, Stanford, CA, USA). The results were visualized as an intensity heatmap with R software. The microarray data have been submitted to the GEO repository: GSE135259.

### RNA-Seq analysis

HEK293FT cells were treated with pcDNA3.1-hISLR-HA or empty vector for 24 hours. RNA was extracted with a standard procedure. RNA quality was assessed on a 2100 Expert Bioanalyzer (Agilent). RNA samples were sent for library preparation and sequencing on the Illumina NovaSeq 6000 platform by Shanghai Majorbio Bio-pharm Technology. The data were analyzed on the free online platform of Majorbio I-Sanger Cloud Platform (www.i-sanger.com) or using R software. The RNA-Seq data have been submitted to the GEO repository: GSE135280.

### Luciferase assay for Islr promoter activity

The sequence for *Islr* is located on chromosome 9, (NC_000075.6, base pairs 58156246..58159236, complement) in the mouse genome. An approximately 2 kb region upstream of transcript start site (TSS) was identified as the *Islr* promoter in this study, which is located at chromosome 9, (NC_000075.6: base pairs C58161222..58159222) and cloned into the pGL3-Basic reporter constructs. Binding sites 1 and 2 of ETS1 are located at base pairs 58159520-58159506 and base pairs 58160294-58160280, respectively. The construct was verified by DNA sequencing. Nucleotides at Ets1 binding sites were mutated by carrying out site-directed mutagenesis (BGI, Shenzhen, China). The wild type and mutant plasmids were transfected with phRL-TK control plasmid to UC-MSCs, respectively. The luciferase activities of firefly and renilla were measured after 24 hours of transfection by using Dual-Glo luciferase assay kit (Promega) according to the manufacturer’s instructions.

### Mass spectrometric analysis

HEK293FT cells were treated with HA-hISLR or empty vector for 24 hours in 10 cm dishes. Lysates were prepared with a standard procedure and immunoprecipitated with an HA antibody and protein G magnetic beads. Immune complexes were resolved on the SDS-PAGE and digested with Trypsin Gold (Mass Spectrometry Grade, Promega) according to the manufacturer’s instructions. Extracted peptides samples were purified and concentrated using ZipTip pipette tips (Millipore Corporation) following the manufacturer’s directions. The peptides eluted from the ZipTip tips were sent for mass spectrometric analysis. Mass spectrometry analysis was performed in a data-dependent manner, with full scans (350–1,600 m/z) acquired using an Orbitrap mass analyzer at a mass resolution of 70,000 at 400 m/z in Q Exactive.

### Chromatin immunoprecipitation (ChIP) assay

A ChIP assay was performed using the SimpleChIP enzymatic chromatin immunoprecipitation kit (Cell Signaling Technology, #9002) according to the manufacturer’s protocol, with modifications. Briefly, harvested UC-MSCs were first crosslinked with 1% (v/v) formaldehyde for 10 min. After isolating the nuclei by lysis of the cytoplasmic fraction, the immunoprecipitation preparations were incubated with micrococcal nuclease for 20 min at 37°C with frequent mixing to digest the DNA into fragments of 150–900 bp. The sonicated nuclear fractions were divided for input control and were incubated with anti-Ets1, anti-H3 (as a positive control), and anti-IgG (as a negative control) at 4°C overnight. The recruited genomic DNA obtained from the ChIP assay was quantified by qRT-PCR with primers specific for the Ets1 binding elements of the *Islr* promoter regions. The primers are used in Appendix table 3.

### Statistical analysis

All analyses were performed at least in triplicate, and the means obtained were subjected to independent *t*-tests. In the figures, asterisks denote statistical significance (\**P* < 0.05; \*\**P* < 0.01; \*\*\**P* < 0.001). All data are reported as the mean ± standard deviation (SD). The means and SDs from at least three independent experiments are presented in all graphs.

## Acknowledgements

ZY is supported by the National Natural Science Foundation of China (No. 81772984, 81572614); the Major Project for Cultivation Technology (2016ZX08008001); Basic Research Program (2015QC0104, 2015TC041, 2016SY001, 2016QC086); SKLB Open Grant (2018SKLAB6-12).

## Author Contributions

ZY designed research; JX, YT, XS, YTian, MD, SD, CL, YS and PL performed research; JX, YLou, YLi, BZ, YC, ZL, MP, QM, ZZ and ZY analyzed data; JX, ZZ and ZY wrote the manuscript.

## Conflict of interest

The authors have declared that no conflict of interest exists.

## Data availability

The datasets and computer code produced in this study are available in the following databases:

RNA-Seq data: Gene Expression Omnibus GSE135259 (https://www.ncbi.nlm.nih.gov/geo/query/acc.cgi?acc=GSE135259)

RNA-Seq data: Gene Expression Omnibus GSE135280 (https://www.ncbi.nlm.nih.gov/geo/query/acc.cgi?acc=GSE135280)

## Expanded View figure legends

**Figure EV1. *Islr* is primarily expressed in stromal cells and upregulated in colitis and colorectal cancer.**

(A) *In situ* hybridization for *Islr* with RNAscope probe in mouse colons from mouse embryos at E16.5, young mice at postnatal day 10 and adult mice at 8 weeks old. *Ppib*, positive control. *DapB*, negative control. Scale bar: 25 μm. (B) Western blotting for Islr in colonic tissues from mice at indicated timepoints upon DSS treatment and after DSS removal. GAPDH was used as a loading control. (B) Immunohistochemistry for ISLR in human colorectal adenocarcinoma at indicated pathological stages. Scale bar: 50 μm.

**Figure EV2. No apparent phenotype observed in *Islr* cKO mice at homeostasis.** (A) X-gal staining showing the *Twist2^+/Cre^*activity in stromal cells using *Twist2^+/Cre^;R26R^LacZ^*mice. Scale bar: 50 μm. (B) Schematics of Islr genomic locus and strategy for generating loxP targeted alleles. (C) qRT-PCR for *Islr* in intestinal and colonic tissues from control and cKO mice. \*\**P* < 0.01. (D) Western blotting for Islr in colonic tissues from control and cKO mice. α-Tubulin Was used as a loading control. (E) *In situ* hybridization for *Islr* with RNAscope probe in colons from control and cKO mice, showing that *Islr* is deleted in stromal cells, while not in the epithelial cells. Dashed lines marked the border of epithelium. Scale bar: 25 μm. (F) H&E staining of colons from control and cKO mice. Scale bar: 100 μm. (G) Double immunofluorescence for Ki67 and β-catenin, immunohistochemistry for cleaved caspase3 (Casp3), and PAS staining in colons from control and cKO mice. n=3. Scale bar: 50 μm. (H) Quantification for crypt number, Ki67^+^ cells, Casp3^+^ and PAS^+^ cells in Panel G. (I) Immunohistochemistry for CD45 and immunofluorescence for F4/80 in colons from control and cKO mice. n=3. Scale bar: 100 μm.

In C, data are presented as mean ± SD. \*\**P* < 0.01 (Student’s t-test).

**Figure EV3. Deletion of *Islr* in stromal cells impaired intestinal epithelial regeneration in TNBS-induced colitis.**

(A) Quantification of body weight change in control (n=8) and cKO (n=8) mice after TNBS treatment. \*\**P* < 0.01; \*\*\**P* < 0.001. (B) Gross images of colons from control (n=5) and cKO (n=5) mice four days after TNBS treatment, and quantification of colon length from control and cKO mice. \*\*\**P* < 0.001. (C) Histological images of colonic inflamed mucosa from control and cKO mice at indicated timepoints after TNBS treatment. Scale bar: 100 μm. (D) Quantification clinical score of inflamed mucosa in control and cKO mice. \*\*\**P* < 0.001. (E) Double immunofluorescence for Ki67 and b-catenin in colonic inflamed mucosa from control and cKO mice at indicated timepoints after TNBS treatment. Scale bar: 100 μm.

In A, B and D, data are presented as mean ± SD. \*\**P* < 0.01; \*\*\**P* < 0.001 (Student’s t-test).

**Figure EV4. Deletion of *Islr* in stromal cells suppressed epithelial Yap signaling.**

(A) Heatmaps of differentially expressed genes (DEGs) in colons from control and cKO mice one day after DSS removal. Note: the cutoff is *P* < 0.05 and fold > 2. (B) KEGG pathway analysis of DEGs in cKO mice. (C) RT-PCR analysis for *CTGF*, *ANKRD1*, *CYR61* and *FSTL1* in NCM460 colon epithelial cells transfected with vector or YAP1-5SA. \**P* < 0.05; \*\*\**P* < 0.001.

In C, data are presented as mean ± SD. \**P* < 0.05; \*\**P* < 0.01; \*\*\**P* < 0.001 (Student’s t-test).

**Figure EV5. Stromal cell-secreted Islr suppressed epithelial Hippo signaling and activated YAP activity.**

(A) KEGG enrichment analysis in DEGs of hISLR overexpressing HEK293FT cells. (B) Western blotting for HA and ISLR in the supernatant from NCM460 colon epithelial cells transfected with pcDNA3.1 or pcDNA3.1-hISLR-HA plasmids. (C) Western blotting for pMST1/2, MST1, pMOB1 and MOB1, pYAP1 and YAP1 in lysates from NCM460 colon epithelial cells cultured in the supernatants with hISLR-HA. β-actin was used as a loading control. (D) Western blotting for HA and ISLR in the supernatant from HCT116 colorectal cancer cells transfected with pcDNA3.1 or pcDNA3.1-hISLR-HA plasmids. (E) Western blotting for pMST1/2, MST1, pMOB1 and MOB1, pYAP1 and YAP1 in lysates from HCT116 colorectal cancer cells cultured in the supernatants with hISLR-HA. β-actin was used as a loading control. (F) Western blotting for YAP in nuclear proteins isolated from NCM460 colon epithelial cells cultured in the supernatant with or without hISLR-HA. Histone H3, a positive control for nuclear proteins. α-Tubulin, a positive control for cytoplasmic proteins. (G) The schematics showing that eleven FAT1 segments in the Cadherin domain were cloned into the Flag-tagged expression vector.

